# Song familiarity relies on evidence accumulation

**DOI:** 10.1101/2025.08.25.672171

**Authors:** Jared R. Girard, Aaron Bishop, Cameron D. Hassall

## Abstract

Familiarity judgements are thought to involve *evidence accumulation*, a decision-making process in which information is gathered over time until a threshold is reached. Previous work has identified a scalp-recorded signature of evidence accumulation called the central-parietal positivity (CPP). We built on this previous work and recorded electroencephalography (EEG) while participants listened to several melodies, instructing participants to respond as soon as the song felt familiar. A prominent CPP was noted, time-locked to decisions. We then used linear regression to unmix overlapping neural responses and observed a stimulus-locked indicator of evidence accumulation that increased in amplitude just prior to a familiarity decision. This result suggests that song familiarity relies on evidence accumulation, with individual notes in a familiar song acting as “evidence”.

## Song familiarity relies on evidence accumulation

Familiarity is the feeling that we’ve encountered a stimulus previously and is often contrasted with recall, the retrieval of specific information from a past event. The relationship between familiarity and recall has been debated, but both processes seem to be important in recognition memory. According to some accounts, recognition judgements rely on both the “qualitative” recollection of details and the “quantitative” feeling of familiarity (Yonelinas, 2002). Familiarity often occurs in the absence of recall and “recognition without identification” has been observed for words (Cleary & Greene, 2000; Peynircioǧlu, 1990), images (Curran & Cleary, 2003), and songs (Kostic & Cleary, 2009).

One possible mechanism underlying familiarity judgements is *evidence accumulation*, a decision-making process by which information is gathered over time until a threshold is reached. Here, familiarity is viewed as a graded signal which, if large enough to exceed a criterion, leads to a familiarity judgement (e.g., a “know” response in a remember/know task: Tulving, 1985). In other words, the feeling of familiarity builds over time rather than happening instantaneously. Evidence accumulation models are found in many areas of cognitive psychology, including perception, learning, language, value-based decision-making, and memory (Evans & Wagenmakers, 2019). What counts as “evidence” in an evidence accumulation model varies by task and may include observables such as the direction a dot moves in a random dot kinematogram, or unobservables such as information retrieved from memory. For example, according to Ratcliff (1978), memory retrieval works when a stimulus is compared against several items in memory, with each comparison initiating a separate accumulation process. These processes terminate when a criterion is reached, leading to either a “match” or “non-match” outcome.

Previous work has found convincing behavioural evidence that recognition memory relies on evidence accumulation. In particular, evidence accumulation models do an excellent job of explaining patterns in accuracy and response time (Evans & Wagenmakers, 2019; Osth et al., 2018; Ratcliff, 1978). In contrast, neural evidence for evidence accumulation during recognition memory has been mixed. For example, van Vugt and colleagues (2016) recorded electroencephalography (EEG) while participants made old/new judgements in a face recognition task^1^. To identify the neural correlates of evidence accumulation, van Vugt et al. (2016) used regression to locate ramping activity occurring between a memory probe and a participant’s response. They focused their analysis on several brain oscillations, but did not find any ramping signals, which they interpreted as evidence against a role for evidence accumulation in recognition memory. However, van Vugt and colleagues (2019) later identified a ramping stimulus-locked ERP component called the central parietal positivity (CPP), which they interpreted as indicating an evidence accumulation process during recognition memory. Additionally, they related the observed signal to “strength of evidence”, which they defined as the difference between the current face and the remembered face. Their results align with previous work showing that the CPP is a neural marker of *perceptual* evidence accumulation (Devine et al., 2019; Dou et al., 2024; Kelly & O’Connell, 2013; Loughnane et al., 2018; O’Connell et al., 2012; Pisauro et al., 2017; Ruesseler et al., 2022; Twomey et al., 2015).

Prior work by van Vugt et al. (2016; 2019) is conflicting and has two limitations. First, it is unclear whether the central-parietal activity they observed was related to the appearance of the memory probe, the subsequent decision/motor response, or both. Second, they used a delayed match-to-sample task, which may have included both a recollection component and a familiarity component. Whether or not familiarity alone elicits a neural measure of evidence accumulation is currently unknown.

To address these limitations, we recorded EEG while participants completed a song familiarity task. We focused on song familiarity under the assumption that individual notes could be treated as observable evidence in favour of a familiarity decision^2^. Participants listened to song melodies and indicated familiarity with a button press. For familiar songs, participants were asked to identify the song. Thus, although we did not have precise control over the strength of evidence each note provided^3^, we planned to test the effect of “strength of evidence” by contrasting stimulus-locked activity for *unfamiliar* notes and *familiar* notes (i.e., those preceding a familiarity judgement).

To address the issue of overlap between stimulus-locked activity and response-locked activity, we used a regression-based approach called *deconvolution* (Ehinger & Dimigen, 2019; Smith & Kutas, 2015a, 2015b). This allowed us to examine stimulus-locked activity and response-locked activity in isolation. Based on previous work (O’Connell & Kelly, 2021; van Vugt et al., 2019), and if familiarity processes rely on evidence accumulation, one or both of the following results should emerge. First, EEG time-locked to familiar notes should be more positive at central-parietal locations compared to unfamiliar notes (i.e., an enhanced stimulus-locked CPP). Second, EEG time-locked to familiarity decisions should be ramping and positive (i.e., an enhanced response-locked CPP).

## Method

### Participants

No power analysis was conducted because we had no prediction about the amplitude of the CPP (familiarity-related CPPs have not been measured previously). However, prior studies have observed a reliable CPP for other decision types for sample sizes ranging from 18 to 25 participants (Dou et al., 2024; Kelly & O’Connell, 2013; O’Connell et al., 2012; Pisauro et al., 2017; van Vugt et al., 2019). We chose to test 30 participants with an average age of 22.63 years (*SD* = 5.38). Ten of the participants self-reported as male; the rest self-reported as female. Three participants self-reported as left-handed, one as ambidextrous, and the rest were right-handed. Five of the participants had coarse curly hair and were included in the study but required a greater-than-average amount of electrolytic gel to lower electrode impedances. Participants provided informed consent and were compensated for their time with course credit. The study was approved by the Research Ethics Board at MacEwan University. One participant was excluded from the study due to technical issues with the EEG amplifier. Two additional participant were excluded due to low trial numbers (two or fewer familiar songs).

### Apparatus and procedure

Participants sat approximately 625 mm from a 527X296 mm display (60Hz, 1920 by 1080 pixels, Elo 2402L). Visual and auditory stimuli were presented using Psychtoolbox (Brainard, 1997; Pelli, 1997) in MATLAB 2023a (Mathworks). Audio was produced using an audio interface (Focusrite Scarlett Solo 4^th^ Generation) and two studio monitors approximately 750 mm from the participant’s left and right ears (JBL 305P MkII). The audio was adjusted to a comfortable level, and participants were asked to minimize head and eye movements throughout the experiment.

Participants then completed a song familiarity task. At the start of each trial, a song melody would play. The melodies were taken from a previous study (Kostic & Cleary, 2009) and consisted of piano notes only. The melodies ranged in duration from 4.78 s to 17.07 s (M = 9.54, SD = 2.31). Participants were instructed to “Press the SPACEBAR as soon as the song feels familiar, regardless of whether or not you can name it.” While the song played, a small fixation cross was displayed on the screen; participants were instructed to keep their eyes at this location. If the participant pressed the spacebar to indicate that the song felt familiar, the display immediately switched to the text prompt “Name, lyric, source?”. Participants were instructed that this was their cue to identify the name of the song, part of the lyrics, or the “song source” such as a show or movie where they had heard it previously. If participants could not identify anything about the song, they were instructed to leave the prompt blank. The prompt disappeared when the participant pressed the enter key and was replaced by four song titles, including the true song title. Participants were instructed to select the title of the song they had just heard. If they selected the correct song title, the selection turned green, and a checkmark appeared; otherwise, the selection turned red, and an ‘x’ appeared. For trials in which the song was unfamiliar to the participant, the song stopped playing after several seconds and the rest of the trial proceeded the same way (the same text prompt followed by the multiple-choice query). See Figure 1 for a trial overview.

**Figure 1.**
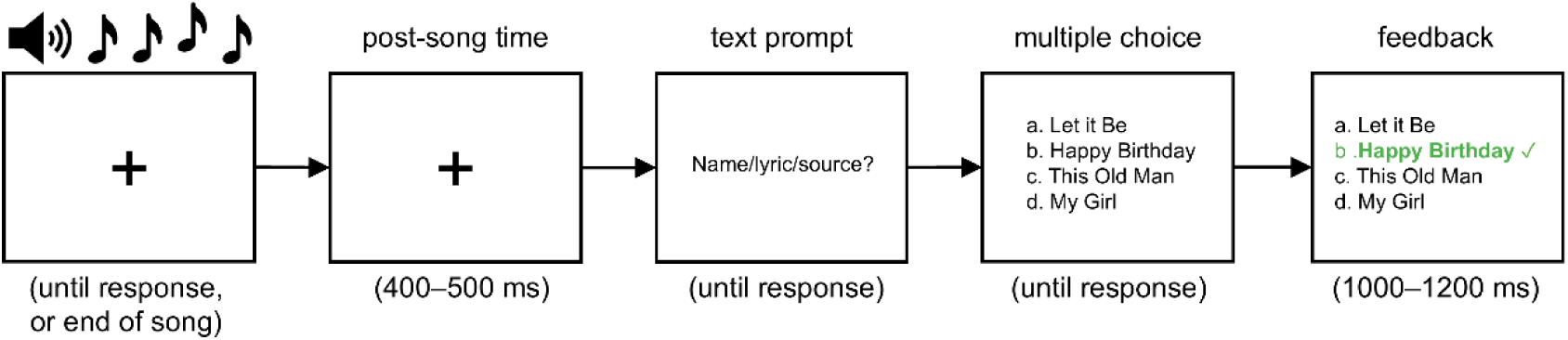
Task details and behavioural data. (a) Participants listened to several songs and made a familiarity judgement (button press for familiar songs, no response for unfamiliar songs). Participants were then given two opportunities to identify the song: a text prompt and a multiple-choice question (with feedback).

### Data collection

For each song, the experimental program recorded the song title, whether the participant responded, the participant’s response to the “Name, lyric, source?” prompt, and the participant’s response to the multiple-choice query. We also recorded 32 electrodes of EEG referenced to electrode Fz using Brain Vision Recorder (Version 1.26.0001, Brain Products GmbH). Thirty electrodes were placed on an elastic cap with standard 10–20 layout (EASYCAP GmbH). Two electrodes were attached to the left and right mastoids to serve as a later reference. Conductive gel was used to lower electrode impedances. The EEG was sampled at 1000 Hz and amplified using a BrainAmp DC (Brain Products GmbH) with a 250 Hz anti-aliasing hardware filter.

## Data analysis

### Behavioural analysis

The behavioural analysis was done in MATLAB 2024b (MathWorks). The goal of the behavioural analysis was to check the feasibility of our event-related EEG analysis. The CPP is similar to another component called the P300, which has been shown to require around 20 trials per condition (Boudewyn et al., 2018; Cohen & Polich, 1997). For each participant and trial, we manually examined what was written in response to the “Name/lyric/source?” prompt. Trials with blank responses were labelled as unidentified, as were trials with statements such as “not sure” or “don’t know”. All other responses were labelled as “identified”, regardless of correctness. For each participant we then calculated the total number of unfamiliar songs and familiar songs. To inform our stimulus-locked analysis, we calculated the total number of *notes* each participant heard for each song type (unfamiliar, familiar). We also calculated the proportion of each song type that could be identified using the text prompt and using the multiple-choice prompt. As mentioned above, we did not check the text response for correctness. However, only correct responses to the multiple-choice query counted as “identified”.

### EEG preprocessing

The EEG was analyzed in MATLAB 2024a (MathWorks) using the EEGLAB library (Delorme & Makeig, 2004). After downsampling to 250 Hz, we applied a bandpass filter (0.1–20 Hz, 60 Hz notch), and rereferenced to the average of the mastoid signals. Ocular artifacts were identified and removed using independent component analysis (ICA). The ICA was trained on the entire recording, minus any samples that were flagged as artifacts. These samples were identified using a sliding-window approach with a 2000 ms window and a step size of 100 ms. If the voltage in the window dropped below -500 µV or exceeded 500 µV, all samples in the window were flagged and excluded from the ICA. Ocular components were then identified using the function *iclabel* and removed from the continuous data set if assigned an “Eye” label with a likelihood exceeding the sum of all “Non-Eye” components (Pion-Tonachini et al., 2019). On average, we removed 3.59 independent components (*SD* = 1.18).

### Event-related analysis

To unmix the stimulus-locked and response-locked signals, we used an approach called deconvolution, which models the ongoing EEG as a linear combination of underlying “regression ERPs” (Burns et al., 2013; Ehinger & Dimigen, 2019; Smith & Kutas, 2015b, 2015a). This approach yields a regression ERP (rERP) for each participant, electrode, and modelled event that could be subjected to the same statistics as an ERP, discussed below. Using a toolbox called Unfold (Ehinger & Dimigen, 2019), we constructed a regression model that included a -1500 to 1500 ms window around unfamiliar notes, familiar notes, and motor responses. After constructing the regression (i.e., after generating the design matrix), we checked each participant’s continuous EEG for artifacts using a 1000 ms sliding window (100 ms step size). If any sample within the window exceeded 75 µV or fell below -75 µV, all samples within the window were flagged as artifacts and excluded from the regression. This was done separately for each electrode; if more than 10% of modelled samples were flagged for a particular electrode then we re-ran the preprocessing step for that participant after removing and interpolating the data from the bad electrode. The data was removed prior to the ICA step and interpolated afterwards. On average we interpolated 0.27 electrodes per participant (*SD* = 0.59). One participant was excluded from further analysis due to excessive artifacts (17 of their electrodes violated our “10% rule”). In total, our EEG analysis consisted of 26 participants.

Regularization was used to prevent model overfitting. We used a first-derivative form of Tikhonov regularization that penalizes the sample-to-sample change in the solution. This imposes a smoothness constraint on the solution (Kristensen et al., 2017; Reichel & Ye, 2008). Ten-fold cross-validation was used to select the optimal regularization parameter for each participant. Error was defined as the average mean-squared error across all electrodes. The following lambda values were tested: 0 (no regularization), 10, 100, 1000, 10,000, 100,000, 1,000,000, 10,000,000, 100,000,000. The optimal lambda for each participant minimized the mean fold error across all ten folds and was consistent across participants: λ = 10,000 (10 participants) or λ = 100,000 (16 participants).

### Parametric regression

The event-related analysis described above has a potential flaw: It treats all note events equally, regardless of their position relative to the familiarity decision. It could be, for example, that the notes immediately preceding a button press are processed differently compared to the notes towards the beginning of the melody. This would align with previous work showing that musical notes are processed in the context of previously heard notes (Sankaran et al., 2024). To test this, we included note position relative to the button press as a parametric regressor in a second regression analysis. The model also included constant (intercept) terms for notes and button presses. Note position was defined as follows: the note immediately preceding the button press was assigned a position of -1, the note before that -2, and so on. We then estimated the rERP for each regressor using the same regularization parameters as before.

### Statistics

To test whether unfamiliar stimulus-locked activity differed from familiar stimulus-locked activity, we used cluster based permutation testing as described by Maris and Oostenveld (2007). For each electrode, we computed a repeated measures t-statistic at each sample point to determine whether the familiar stimulus-locked voltage at that sample point differed from the unfamiliar stimulus-locked voltage. We then identified clusters of sample points for which the t-values fell below the 2.5th percentile or exceeded the 97.5th percentile. Spatial clusters were defined according to a template from the FieldTrip toolbox (Oostenveld et al., 2010). For each spatial cluster, we defined a “cluster mass” as the sum of the absolute values of the temporal cluster t-values; to be included in a cluster mass, the voltage at a sample point had to reach significance for all members of the spatial cluster. We restricted the stimulus-locked analysis to a window from 0 to 600 ms relative to the onset of the note, a typical stimulus-locked ERP time range. To determine whether the observed cluster masses exceeded what could occur by chance, we permuted the participant rERPs by randomly swapping the condition labels on a participant-by-participant basis. This is equivalent to randomly flipping the sign of a data point in a single-sample t-test. We then computed the cluster masses of the permuted waveforms and recorded the maximum cluster mass, or zero if no clusters were found. We tested 1000 permutations in total. Finally, we labelled an observed cluster as “significant” if its cluster mass exceeded 95% of the permuted cluster masses. The reported *p*-value was the proportion of permuted cluster masses exceeding the observed cluster mass. For each significant cluster, we also reported an effect size by averaging the EEG over the cluster electrodes and sample points for each participant and computing:

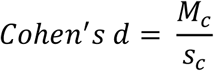

where *Mc* and *sc* represent the mean and standard deviation of the resulting cluster voltages.

A similar procedure was done for the response-locked rERP. Our goal here was to determine whether (and when) the response-locked rERP differed significantly from zero (an “existence test”). We did not split this analysis by note familiarity because responses were only made in the case of a familiar song. To capture decision-related activity, the response-locked analysis was restricted to a window from 1000 ms before the response 100 ms after the response. Permutation testing proceeded as before, except with a single-sample *t*-test (i.e., a repeated-measures test against zero). For both the stimulus-locked and response-locked analysis, and in the case that a significant cluster was found, we visualized the spatial pattern of the signal by calculating the average rERP across the significant temporal clusters for each participant and electrode.

For our parametric regression, we used cluster-based permutation testing to conduct two existence tests. First, we checked to see where/when the note-position rERP differed from zero. If found, a significant cluster would show an effect of note position on stimulus-locked processing. Second, we checked for non-zero response-locked activity, a replication of our previous response-locked analysis. This was necessary because the inclusion of an additional regressor could have influenced the response-locked result.

Finally, we replicated our parametric analysis but excluded identified familiar trials (trials for which the participant could identify the song at the test prompt). This was done to control for the possible effect of recall on stimulus-locked and response-locked activity. In other words, we wanted to be confident that our EEG results were driven by familiarity alone. This analysis required that we exclude two additional participants due to insufficient trial numbers (total N: 24).

## Results

### Behavioural results

Summary statistics, shown in Table 1, show a reasonable number of “familiar” responses. The number of familiar responses ranged greatly across participants, however (Figure 2). We observed that 43% of familiar songs could be identified in writing with a 68% score on the multiple-choice question. We also observed that in all conditions, performance on the multiple-choice question was above chance level (25%). Taken together, our behavioural results suggest that participants completed the task as intended, though there were fewer familiar responses than predicted based on pilot testing.

**Table 1.** Behavioural results showing the mean number of trials and notes, the mean proportion of songs that could be identified, and the mean proportion of songs that were correctly chosen at the final multiple-choice (MC) prompt.

|  | Unfamiliar |  | Familiar |  |
| --- | --- | --- | --- | --- |
|  | *M* | *95% CI* | *M* | *95% CI* |
| Number of songs | 33.8 | [27.8, 39.9] | 38.1 | [30.5, 45.6] |
| Number of notes | 924.6 | [768.0, 1081.1] | 571.1 | [500.7, 641.5] |
| Proportion identified - Text | 0.02 | [0.01, 0.03] | 0.43 | [0.33, 0.52] |
| Proportion identified - MC | 0.42 | [0.36, 0.48] | 0.68 | [0.62, 0.75] |

**Figure 2.**
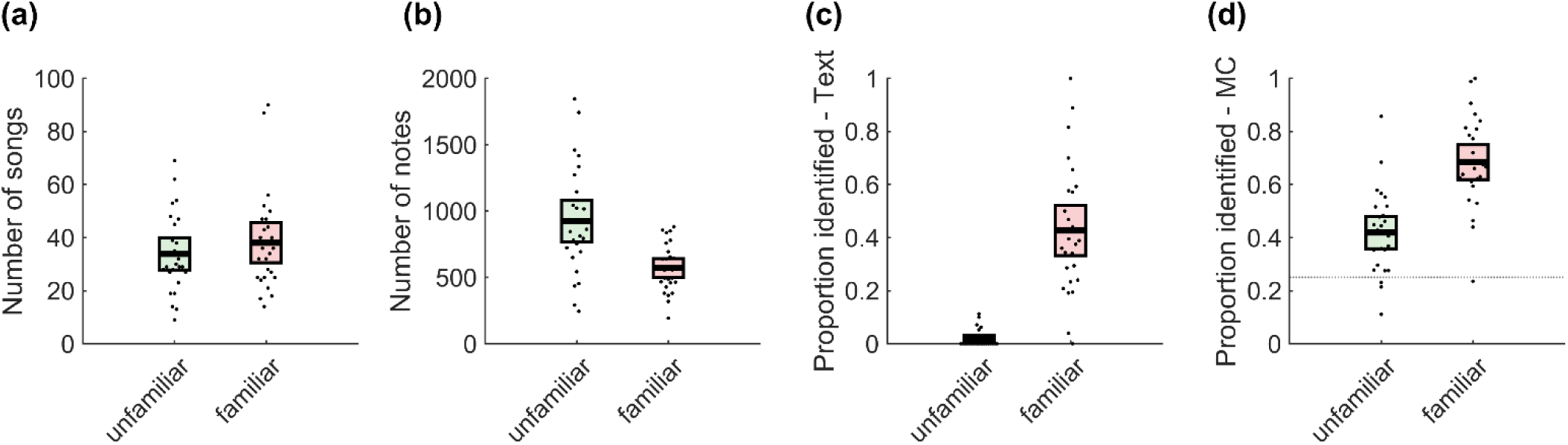
Behavioural results indicate that the task was completed as intended. The number of familiar songs ranged greatly across participants (a). Nevertheless, there were many notes per condition (b). At the text prompt, participants were more likely to identify familiar songs compared to unfamiliar songs (c). Multiple choice (MC) performance was greatest for familiar songs but was above chance level (the dotted line) in both conditions (d). Error bars indicate 95% confidence intervals.

### EEG results

#### Regression ERPs

After using linear regression to model our EEG (Figure 3a) we found no significant clusters differentiating familiar stimulus-locked activity and unfamiliar stimulus-locked activity. We have illustrated the mean stimulus-locked rERP for each condition (familiar, unfamiliar) at two scalp locations: frontal (FCz) and parietal (CPz) – see Figure 3b. This suggests that, on average, stimulus-locked activity did not depend on familiarity (though see our second EEG analysis below). The response-locked result showed activity in line with a CPP, and a significant cluster from -364 ms to 100 ms around O1 was identified (O1, Oz), *p* = .0017, Cohen’s *d* = 0.71 (Figure 3c).

**Figure 3.**
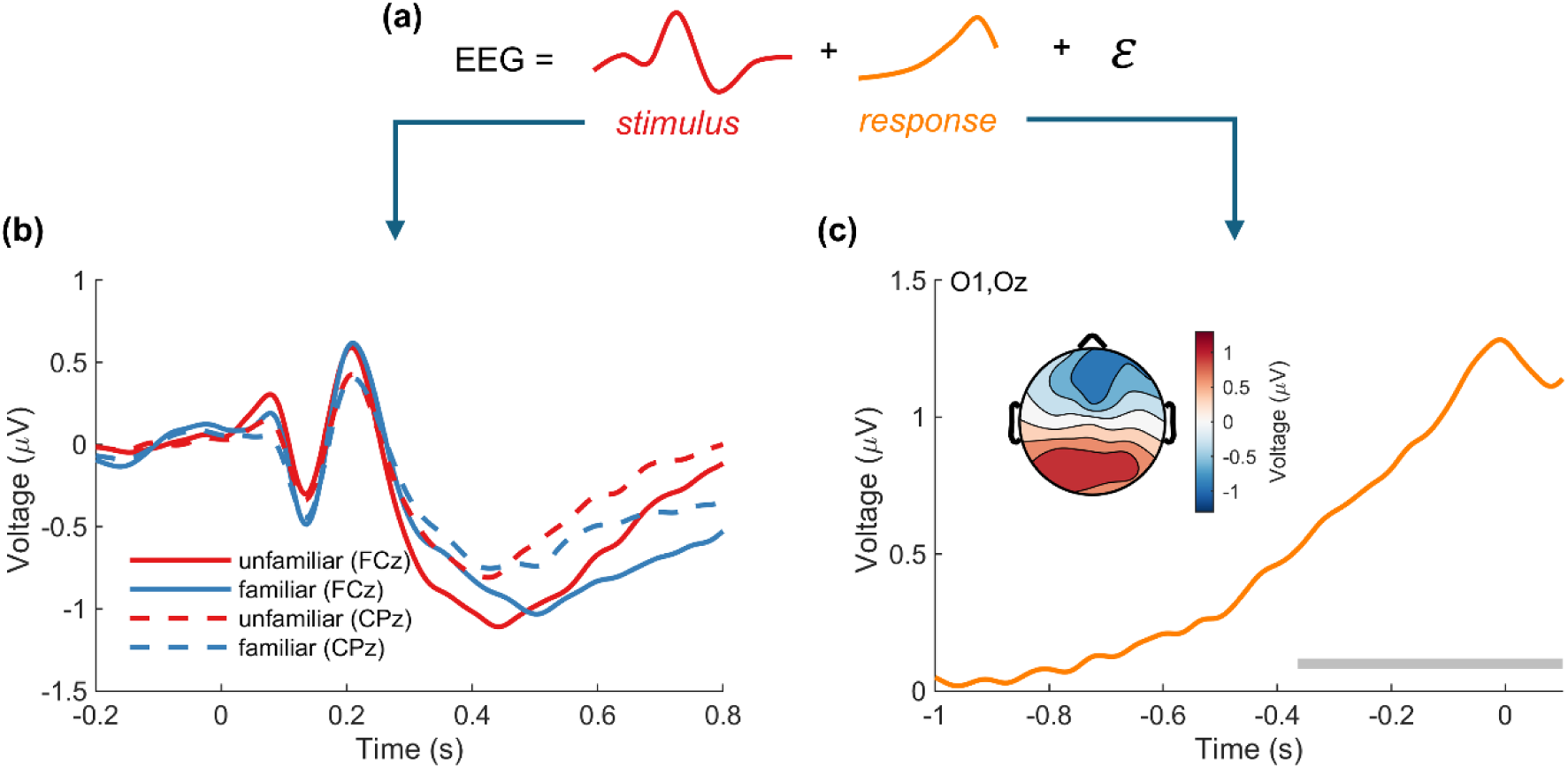
Deconvolution revealed distinct stimulus-locked and response-locked activity. A regression model was constructed that included stimulus-locked activity, response-locked activity, and an error term ε (a). Results showed no significant stimulus-locked differences (b). However, a response-locked CPP was observed (c). The grey bar indicates a significant temporal cluster.

#### Parametric regression

Including note position *p* as a parametric regressor (Figure 4a) split stimulus-locked activity into an intercept component (Figure 4b) and a parametric component (Figure 4c). The parametric results showed a significant stimulus-locked cluster of activity centred at electrode FC1 (FC1, F3, FCz, C3, Cz, Fz) from 308 ms to 600 ms, *p* < .001, Cohen’s *d* = 1.28. Our earlier response-locked result was replicated at a significant cluster from -348 ms to 100 ms centred at O1 (O1, Oz), *p* = .0021, Cohen’s *d* = 0.65. Similar results were obtained near these cluster centres when we excluded identified familiar trials: F4, 468-488 ms, *p* = .018, Cohen’s *d* = 0.66 (stimulus-locked) and O2, -116 ms to 100 ms, *p* = .005, Cohen’s *d* = 0.68 (response-locked).

**Figure 4.**
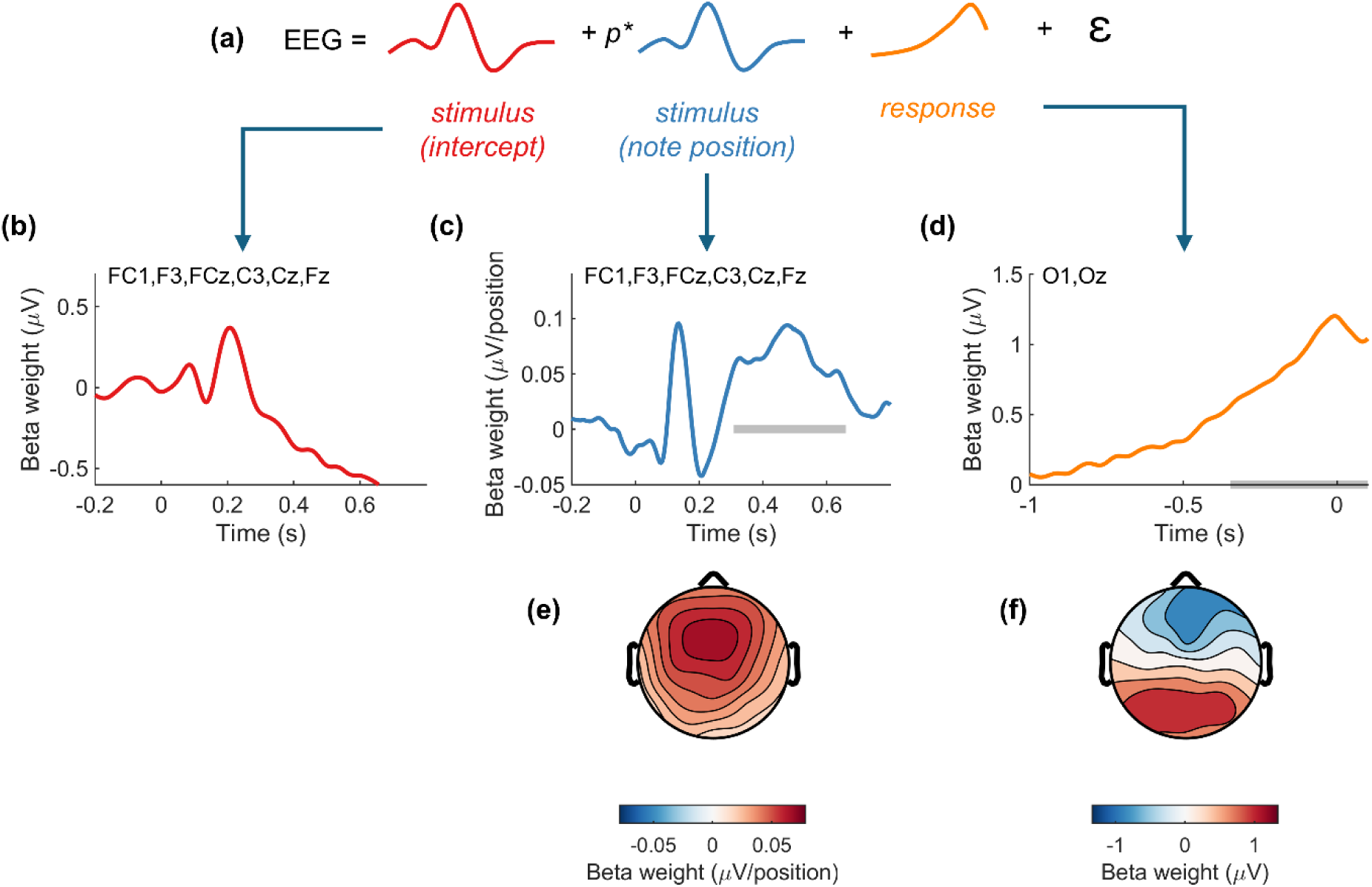
Parametric regression revealed an increasing stimulus-locked positivity leading up to familiarity judgements. Notes preceding button presses were assigned a position *p*, which was then used as a parametric regressor in a regression model that included stimuli (notes), motor responses, and an error term ε (a). When the intercept term (b) and parametric term (c) were estimated, we noted a parametric effect that depended on note position and was maximal at over frontal electrodes (e). We also reproduced our earlier response-locked result (d, f). The grey bars indicate significant temporal clusters.

To visualize the effect of note position on stimulus-locked activity, we reconstructed the EEG using the rERPs. In other words, we combined the intercept rERP result with varying amounts of the parametric rERP result. The reconstruction shows how the stimulus-locked response builds over time (becomes more positive) as the familiarity decision approaches (Figure 5).

**Figure 5.**
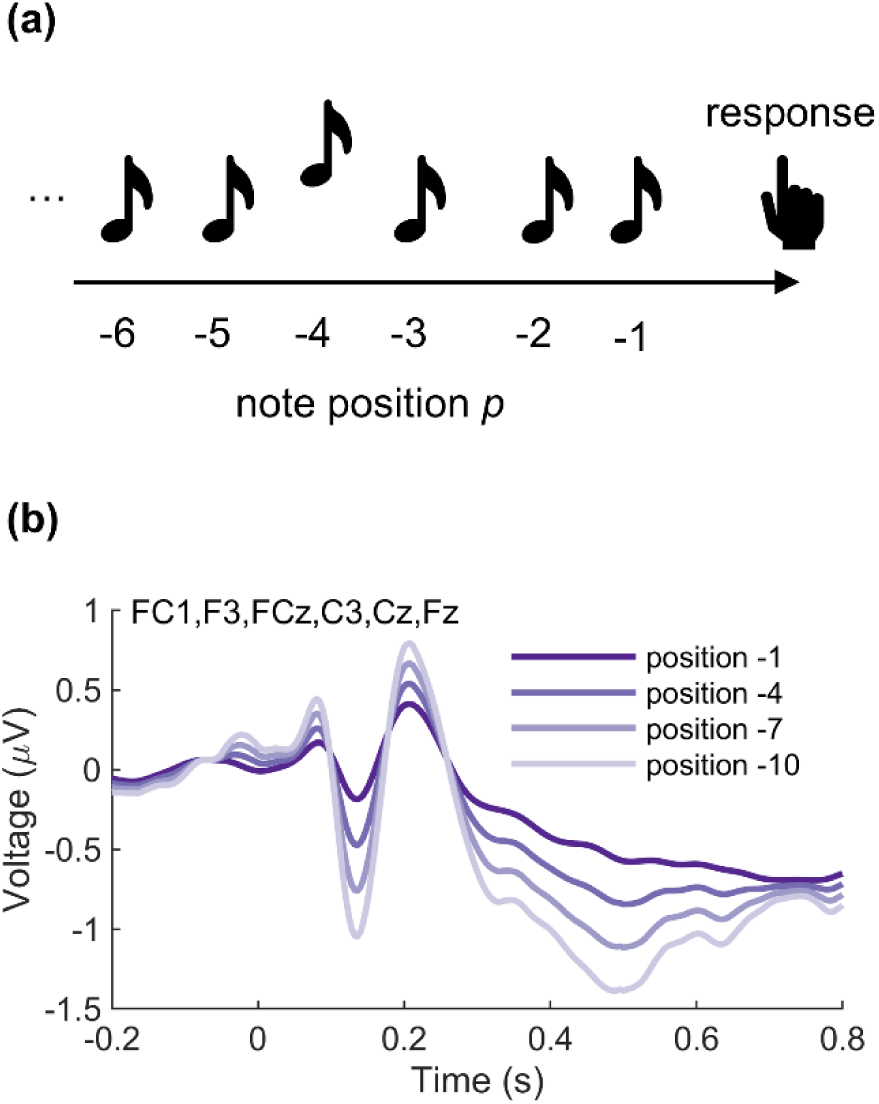
Reconstructed EEG shows the effect of note position on the stimulus-locked signal. As note position draws closer to the button press (a), corresponding stimulus-locked activity increases in amplitude (b).

Out of interest in the difference between *unidentified* familiar trials and *identified* familiar trials, we conducted an exploratory analysis in which we repeated our parametric analysis but modelled these trials separately. We excluded any participant with five or fewer trials in either condition and calculated the mean voltage for each participant and condition at the previously identified clusters (total *N*: 18). Besides being exploratory, this analysis had the additional caveat of low trial numbers, as discussed earlier – many participants had fewer than 20 trials in a condition. We then conducted paired-samples *t*-tests comparing identified familiar and unidentified familiar trials. We found no difference in the note-position effect *t*(17) = 0.22, *p* = .83, Cohen’s *d* = 0.05 and a small difference in the response-locked effect, *t*(17) = 2.13, *p* = .048, Cohen’s *d* = 0.50 (Figure 6).

**Figure 6.**
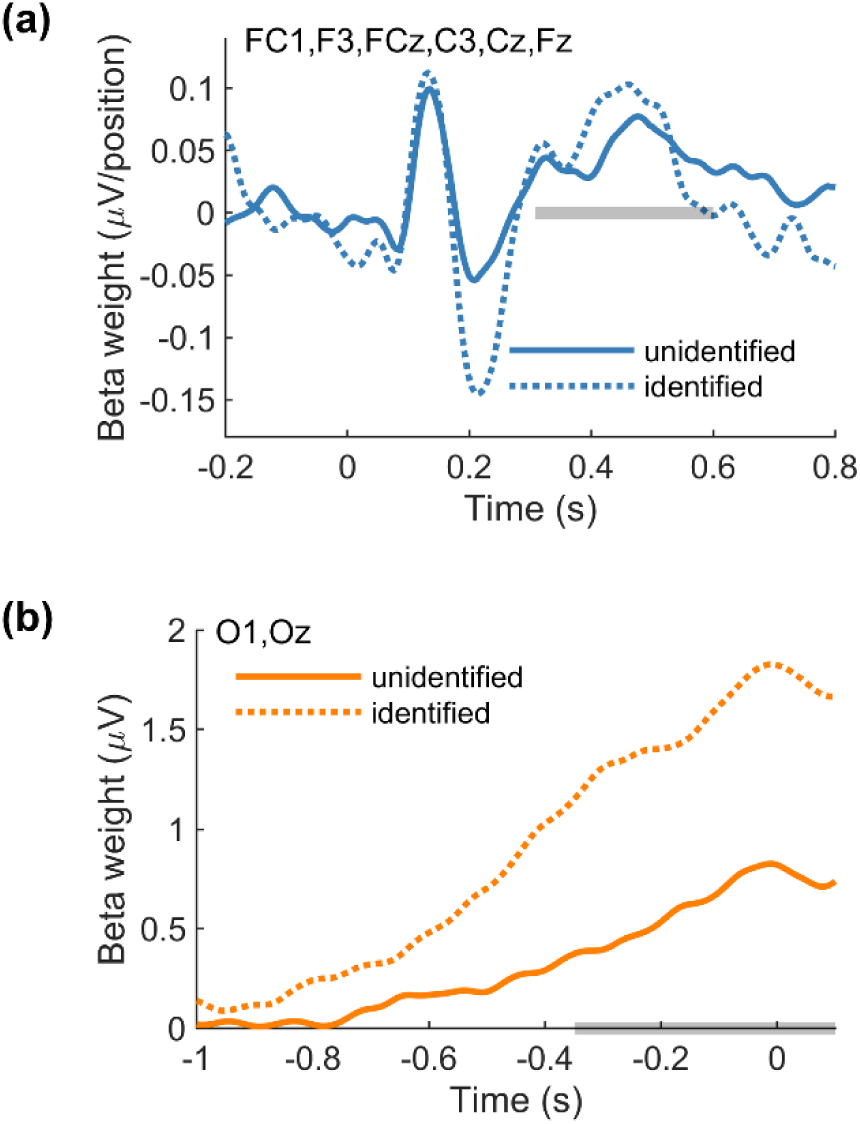
Exploratory parametric analysis, split by song identification. Stimulus-locked activity did not differ between unidentified and identified songs. However, there was a small effect of song identification on the response-locked signal (b). The grey bars indicate the temporal clusters identified in our previous analysis.

## Discussion

Our goal was to build on a small body of previous work showing neural evidence that recognition judgements rely on evidence accumulation. We expanded on this work in two ways. First, we used an unmixing procedure to isolate stimulus-locked and response-locked activity. This allowed us to examine the effect of “strength of evidence” on the stimulus-locked signal independent of response-locked activity. Second, we focused on familiarity judgements as opposed to recall judgements. In line with previous work (Evans & Wagenmakers, 2019; Osth et al., 2018; Ratcliff, 1978; van Vugt et al., 2019), our results show that familiarity judgements rely on evidence accumulation, revealing separate stimulus-locked and response-locked accumulation signals.

In our first EEG analysis, we examined both stimulus-locked and response-locked activity for neural signals related to evidence accumulation. A prominent response-locked CPP was observed, similar to response-locked activity that has been seen previously in perceptual tasks and linked to evidence accumulation (Devine et al., 2019; Dou et al., 2024; Kelly & O’Connell, 2013; Loughnane et al., 2018; O’Connell et al., 2012; Pisauro et al., 2017; Ruesseler et al., 2022; Twomey et al., 2015). However, we found no evidence of stimulus-locked evidence accumulation (i.e., no difference between familiar and unfamiliar notes). This contrasts with previous work showing a stimulus-locked CPP during recall (van Vugt et al., 2019). We note two important differences between previous work and our work. First, our task emphasized familiarity, not recall. Second, we used an unmixing procedure to isolate stimulus-locked and response-locked activity. The reason why unmixing is necessary is to rule out the possibility that the observed activity is artefactual and the result of overlap between adjacent events (Frömer et al., 2024). Thus, our first EEG analysis suggests that familiarity decisions (but not familiar stimuli) elicit a CPP.

Our first EEG analysis examined *average* stimulus-locked activity; our second analysis instead tested whether stimulus-locked activity changes over time. By including note position as a parametric regressor we found evidence of a positive-going change at frontal electrode sites: more positive for notes closer in time to the familiarity decision. This effect had a frontal scalp distribution that was distinct from the CPP observed at the time of the response. Frontal cortex is known to be involved in evidence accumulation during perceptual decisions (Brosnan et al., 2020; Rahnev et al., 2016) and frontal stimulus-locked evidence accumulation signals have been observed during interval timing tasks (Ofir & Landau, 2022) and perceptual judgement tasks (Gherman et al., 2024). We interpret our parametric result as evidence of a stimulus-locked contribution to evidence accumulation.

The precise mechanism behind our stimulus-locked effect is unclear and possibly beyond the scope of this project. Some have argued that frontal stimulus-locked activity in the presence of a parietal evidence-accumulation signal may simply reflect increased attention to decision-relevant stimuli (Gherman et al., 2024). In line with this perspective, our stimulus-locked effect had a timing and topography consistent with the P300, an ERP component linked with several cognitive processes including enhanced arousal to targets in a target detection task (Nieuwenhuis et al., 2005). Under this view, the attention-related activity would be incidental and driven by an accumulation process nearing a decision threshold.

Others have argued that frontal signals such as error- and/or conflict-related activity might *contribute* to the accumulation process, e.g. as an internal evidence signal (Desender et al., 2021; Murphy et al., 2015; Stone et al., 2022, 2024; Wendelken et al., 2009). In line with this perspective, the observed effects are consistent with an ERP component called the N400, which is related to the processing of meaningful stimuli (including tones: Kutas & Federmeier, 2011). A key driver of the N400 and a key feature of music familiarity is the prediction of upcoming notes based on the preceding context (Leaver et al., 2009; Malekmohammadi et al., 2023). Although our task was not designed to elicit the N400 (i.e., we did not manipulate the notes), we speculate that note predictability – as indexed by the N400 – could have served as internal evidence in favour of a familiarity decision. Put another way: A song might be judged as familiar if the notes start to feel predictable. The link between familiarity and prediction has been studied previously. For example, participants are more likely to report a feeling of prediction (i.e., of knowing what happens next) when viewing a familiar video compared to an unfamiliar video (Cleary et al., 2021). In general, predictions about the future rely on remembering the past (Schacter et al., 2007).

Our task was not designed to differentiate between “attention” and “prediction” explanations of our stimulus-locked effect. Future work could manipulate attention using dual tasking or masking. By introducing unexpected (i.e., altered) notes, it might also be possible to examine the effect of stimulus predictability on familiarity decisions. These manipulations could help determine whether frontal activity contributes to evidence accumulation or is merely driven by it. This brings up another limitation of the present work. We are claiming here that song familiarity relies on evidence accumulation because we identified a response-locked CPP. This is a reverse inference – inferring a specific cognitive process from an observed signal – the validity of which relies on the specificity of the observation (Poldrack, 2006). Our confidence that the observed ramping activity is related to evidence accumulation as opposed to some other cognitive process comes from a careful reading of previous CPP work (Devine et al., 2019; Dou et al., 2024; Kelly & O’Connell, 2013; Loughnane et al., 2018; O’Connell et al., 2012; Pisauro et al., 2017; Ruesseler et al., 2022; Twomey et al., 2015) and the fact that we know of no other central-parietal positive-going decision-locked signals. This previous work is high in internal validity but – we would argue – less ecologically valid compared to our task. Part of our motivation for the current work was to apply what has been learned previously about the CPP to a naturalistic task where measuring decision variables precisely is unfeasible.

Taken together, our EEG results suggest that song familiarity relies on evidence accumulation. This aligns with previous work in which evidence accumulation models have been applied successfully to recognition memory (Criss, 2010; Ratcliff et al., 2004; Starns et al., 2012). It also aligns with work in which these models have been applied specifically to familiarity (Cox & Shiffrin, 2012, 2017; Gomilsek et al., 2025; Pan & Hu, 2025; Weigard et al., 2024). Furthermore, our exploratory analysis suggests that familiarity plus recollection may elicit greater accumulation-related activity compared to familiarity alone. This difference is not apparent in the stimulus-locked signal, but rather around the time of the familiarity decision as an amplitude increase (and possible an increase in slope as well – see Figure 6b). Similar response-locked differences are seen when comparing easy/difficult decisions in the domains of perception (Dou et al., 2024; Kelly & O’Connell, 2013; O’Connell et al., 2012; Ruesseler et al., 2022), value (Pisauro et al., 2017), and memory (van Vugt et al., 2019). These differences have been interpreted as representing different rates of evidence accumulation (the “drift rate”) – steeper for “easy” decisions, shallower for “hard” decisions. In our case, this interpretation would suggest faster accumulation when familiarity is followed by recall than by familiarity alone. The enhanced accumulation signal, when viewed as “strength of familiarity”, is consistent with work showing that familiarity can improve source memory (Duarte et al., 2006; Hicks et al., 2002; Mollison & Curran, 2012; Wais et al., 2008).

Many decisions are made gradually, not suddenly, and can be explained by carefully examining both behaviour and neural activity in highly controlled laboratory experiments. We expanded this work to a naturalistic context and found neural evidence that familiarity decisions rely on evidence accumulation. Our work highlights both the challenge and the value of moving towards an ecologically valid understanding of the neural basis of decision-making in memory and other domains.

## Author Note

This research was funded by a Natural Sciences and Engineering Research Council of Canada (NSERC) Discovery Grant to Cameron D. Hassall (RGPIN 2024-04848).

The authors have no conflict of interest.

## Author Contributions

**Jared R. Girard:** Investigation; writing – review and editing. **Aaron Bishop:** Investigation; writing – review and editing. **Cameron D. Hassall:** Conceptualization; formal analysis; investigation; data curation; writing – original draft preparation; writing – review and editing; supervision.

## Data Availability Statement

EEG dataset is available at https://doi.org/10.18112/openneuro.ds005876.v1.0.1. Analysis scripts are available at https://github.com/chassall/songfamiliarity.

## Footnotes

1 EEG is particularly well-suited to studying evidence accumulation due to its temporal resolution (the accumulation process happens on the order of seconds, or less).

2 In addition to individual notes, song familiarity also depends on rhythm (Kostic & Cleary, 2009) and timbre (Faubion-Trejo & Mantell, 2022).

3 In contrast, studies examining perceptual decisions often have precise control over the information presented to participants, which helps when interpreting the CPP. For example, random dot kinematograms are commonly used to study evidence accumulation by carefully controlling the simultaneous motion of many dots (Dou et al., 2024; Kelly & O’Connell, 2013; Kohl et al., 2020; Ruesseler et al., 2022). The participant’s task is to report on the overall direction of motion by integrating over all the dots. Each change in the overall coherence of the dots can be viewed as a piece of evidence in favour of a particular decision, the magnitude of which can be directly linked to the slope and/or amplitude of the CPP (Kelly & O’Connell, 2013).

